# Leveraging Existing 16S rRNA Gene Surveys to Identify Reproducible Biomarkers in Individuals with Colorectal Tumors

**DOI:** 10.1101/285486

**Authors:** Marc A Sze, Patrick D Schloss

**Author notes:** Co-author.

## Abstract

An increasing body of literature suggests that both individual and collections of bacteria are associated with the progression of colorectal cancer. As the number of studies investigating these associations increases and the number of subjects in each study increases, a meta-analysis to identify the associations that are the most predictive of disease progression is warranted. We analyzed previously published 16S rRNA gene sequencing data collected from feces and colon tissue. We quantified the odds ratios (ORs) for individual bacterial taxa that were associated with an individual having tumors relative to a normal colon. Among the fecal samples, there were no taxa that had significant ORs associated with adenoma and there were 8 taxa with significant ORs associated with carcinoma. Similarly, among the tissue samples, there were no taxa that had a significant OR associated with adenoma and there were 3 taxa with significant ORs associated with carcinoma. Among the significant ORs, the association between individual taxa and tumor diagnosis was equal or below 7.11. Because individual taxa had limited association with tumor diagnosis, we trained Random Forest classification models using only the taxa that had significant ORs, using the entire collection of taxa found in each study, and using operational taxonomic units defined based on a 97% similarity threshold. All training approaches yielded similar classification success as measured using the Area Under the Curve. The ability to correctly classify individuals with adenomas was poor and the ability to classify individuals with carcinomas was considerably better using sequences from fecal or tissue.

**Importance:** Colorectal cancer is a significant and growing health problem in which animal models and epidemiological data suggest that the colonic microbiota have a role in tumorigenesis. These observations indicate that the colonic microbiota is a reservoir of biomarkers that may improve our ability to detect colonic tumors using non-invasive approaches. This meta-analysis identifies and validates a set of 8 bacterial taxa that can be used within a Random Forest modeling framework to differentiate individuals as having normal colons or carcinomas. When models trained using one dataset were tested on other datasets, the models performed well. These results lend support to the use of fecal biomarkers for the detection of tumors. Furthermore, these biomarkers are plausible candidates for further mechanistic studies into the role of the gut microbiota in tumorigenesis.

## Background

Colorectal cancer (CRC) is a growing world-wide health problem in which the microbiota has been hypothesized to have a role in disease progression (1, 2). Numerous studies using murine models of CRC have shown the importance of both individual microbes (3–7) and the overall community (8–10) in tumorigenesis. Numerous case-control studies have characterized the microbiota of individuals with colonic adenomas and carcinomas in an attempt to identify biomarkers of disease progression (6, 11–17). Because current CRC screening recommendations are poorly adhered to due to a person’s socioeconomic status, test invasiveness, and frequency of tests, development and validation of microbiota-associated biomarkers for CRC progression could further attempts to develop non-invasive diagnostics (18).

Recently, there has been an intense focus on identifying microbiota-based biomarkers yielding a seemingly endless number of candidate taxa. Some studies point towards mouth-associated genera such as *Fusobacterium*, *Peptostreptococcus*, *Parvimonas*, and *Porphyromonas* that are enriched in people with carcinomas (6, 11–17). Other studies have identified members of *Akkermansia*, *Bacteroides*, *Enterococcus*, *Escherichia*, *Klebsiella*, *Mogibacterium*, *Streptococcus*, and *Providencia* (13–15). Additionally, *Roseburia* has been found in some studies to be more abundant in people with tumors but in other studies it has been found to be less abundant than what is found in subjects with normal colons (14, 17, 19, 20). There is support from mechanistic studies using tissue culture and murine models that *Fusobacterium nucleatum*, pks-positive strains of *Escherichia coli*, *Streptococcus gallolyticus*, and an entertoxin-producing strain of *Bacteroides fragilis* are important in tumorigenesis (5, 14, 21–24). These results point to a causative role for the microbiota in tumorigenesis as well as their potential as diagnostic biomarkers.

Most studies have focused on identifying biomarkers in patients with carcinomas but there is a clinical need to identify biomarkers associated with adenomas to facilitate early detection of the tumors. Studies focusing on broad scale community metrics have found that measures such as the total number of taxa (i.e. richness) are lower in those with adenomas versus controls (25). Other studies have identified *Acidovorax*, *Bilophila*, *Cloacibacterium*, *Desulfovibrio*, *Helicobacter*, *Lactobacillus*, *Lactococcus*, *Mogibacterium*, and *Pseudomonas* to be enriched in those with adenomas (25–27). The ability to classify individuals as having normal colons or adenomas based solely on the taxa within fecal samples has been limited. However, when 16S rRNA gene sequence data was combined with the results of a fecal immunochemical test (FIT), the ability to diagnose individuals with adenomas was improved relative to using the FIT results alone (12).

A recent meta-analysis found that 16S rRNA gene sequences from members of *Akkermansia*, *Fusobacterium*, and *Parvimonas* were fecal biomarkers for the presence of carcinomas (28). Contrary to previous studies, they found sequences similar to members of *Lactobacillus* and *Ruminococcus* to be enriched in patients with adenoma or carcinoma relative to those with normal colons (12, 15, 16). In addition, they found that 16S rRNA gene sequences from members of *Haemophilus*, *Methanosphaera*, *Prevotella*, and *Succinovibrio* were enriched in patients with adenomas and *Pantoea* were enriched in patients with carcinomas. Although this meta-analysis was helpful for distilling a large number of possible biomarkers, the aggregate number of samples included in the analysis (n=509) was smaller than several larger case-control studies that have since been published (12, 27)

Here we provide an updated meta-analysis using 16S rRNA gene sequence data from both feces (n=1737) and colon tissue (492 samples from 350 individuals) from 14 studies (11–17, 19, 20, 23, 25–27, 29) [Table 1 & 2]. We expand both the breadth and scope of the previous meta-analysis to investigate whether biomarkers describing the bacterial community or specific members of the community can more accurately classify patients as having adenoma or carcinoma. Our results suggest that the bacterial community changes as disease severity worsens and that a subset of the microbial community can be used to diagnose the presence of carcinoma.

**Table 1:** Characteristics of the datasets included in the fecal-based analysis.

| Study | Data Stored | Region | Control (n) | Adenoma (n) | Carcinoma (n) |
| --- | --- | --- | --- | --- | --- |
| Ahn | DBGap | V3-4 | 148 | 0 | 62 |
| Baxter | SRA | V4 | 172 | 198 | 120 |
| Brim | SRA | V1-3 | 6 | 6 | 0 |
| Flemer | Author | V3-4 | 37 | 0 | 43 |
| Hale | Author | V3-5 | 473 | 214 | 17 |
| Wang | SRA | V3 | 56 | 0 | 46 |
| Weir | Author | V4 | 4 | 0 | 7 |
| Zeller | SRA | V4 | 50 | 37 | 41 |

**Table 2:** Characteristics of the datasets included in the tissue-based analyses.

| Study | Data Stored | Region | Control (n) | Adenoma (n) | Carcinoma (n) |
| --- | --- | --- | --- | --- | --- |
| Burns | SRA | V5-6 | 18 | 0 | 16 |
| Chen | SRA | V1-3 | 9 | 0 | 9 |
| Dejea | SRA | V3-5 | 31 | 0 | 32 |
| Flemer | Author | V3-4 | 103 | 37 | 94 |
| Geng | SRA | V1-2 | 16 | 0 | 16 |
| Lu | SRA | V3-4 | 20 | 20 | 0 |
| Sanapareddy | Author | V1-2 | 38 | 0 | 33 |

## Results

### Lower bacterial diversity is associated with higher odds ratio (OR) of tumors

We first assessed whether variation in broad community metrics like total number of operational taxonomic units (OTUs) (i.e. richness), the evenness of their abundance, and the overall diversity of the communities were associated with disease stage after controlling for study and variable region differences. In fecal samples, both evenness and diversity were significantly lower in successive disease severity categories (P-value=0.025 and P-value=0.043, respectively) [Figure 1]; there was no significant difference for richness (P-value=0.21). We next tested whether the lower value of these community metrics translated into significant ORs for having an adenoma or carcinoma. For fecal samples, the ORs for richness were not significantly greater than 1.0 for adenoma or carcinoma (P-value=0.40) [Figure 2A]. The ORs for evenness were significantly higher than 1.0 for adenoma (OR=1.3 (95% Confidence Interval: 1.02 - 1.65), P-value=0.035) and carcinoma (OR=1.66 (1.2 - 2.3), P-value=0.0021) [Figure 2B]. The ORs for diversity were only significantly greater than 1.0 for carcinoma (OR=1.61 (1.14 - 2.28), P-value=0.0069), but not for adenoma (P-value=0.11) [Figure 2C]. Although these ORs are significantly greater than 1.0, it is doubtful that they are clinically meaningful.

**Figure 1:**
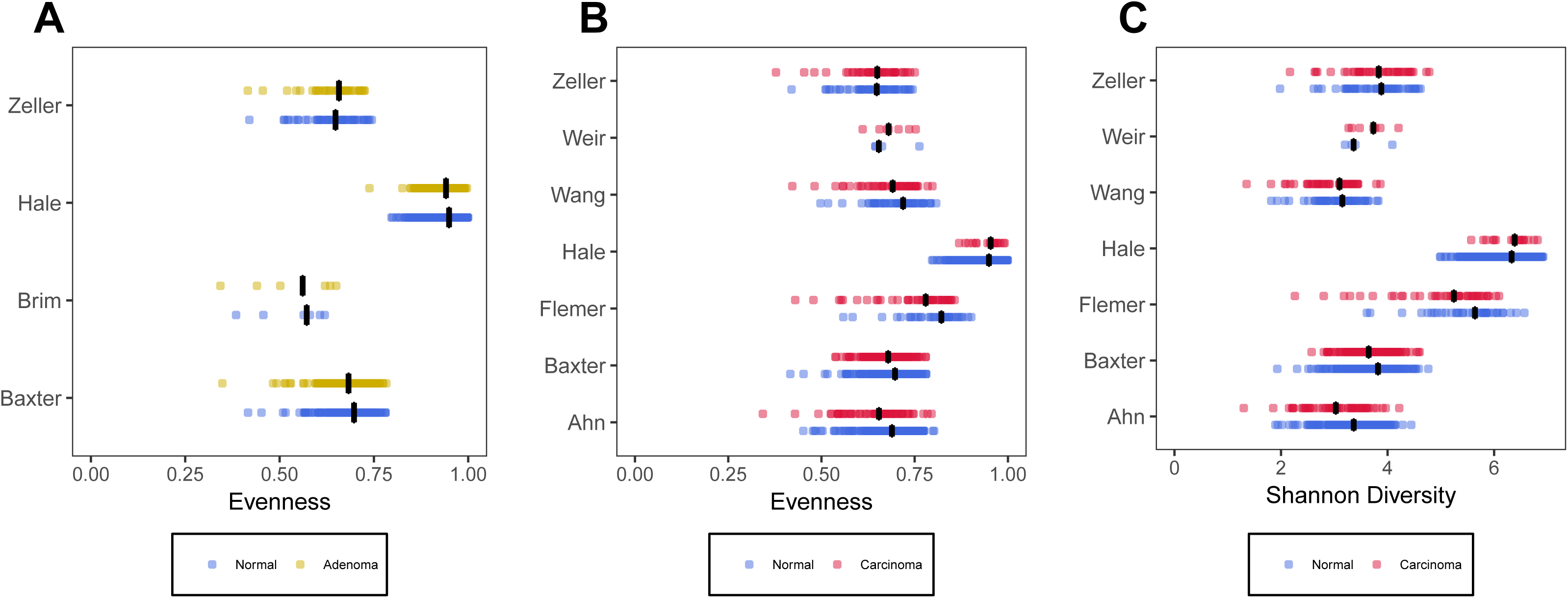
Comparison of alpha diversity indices that were significant between individuals with normal colons, and those with adenomas or carcinomas using data collected from fecal samples. A) Comparison of evenness between individuals with normal colons and adenomas. B) Comparison of evenness between individuals with normal colons and carcinomas. C) Comparison of Shannon diversity between individuals with normal colons and carcinomas. Blue points represent individuals with normal colons and red points represent individuals with either adenomas (panel A) or carcinomas (panel B and C). The black lines represent the median value for each group.

**Figure 2:**
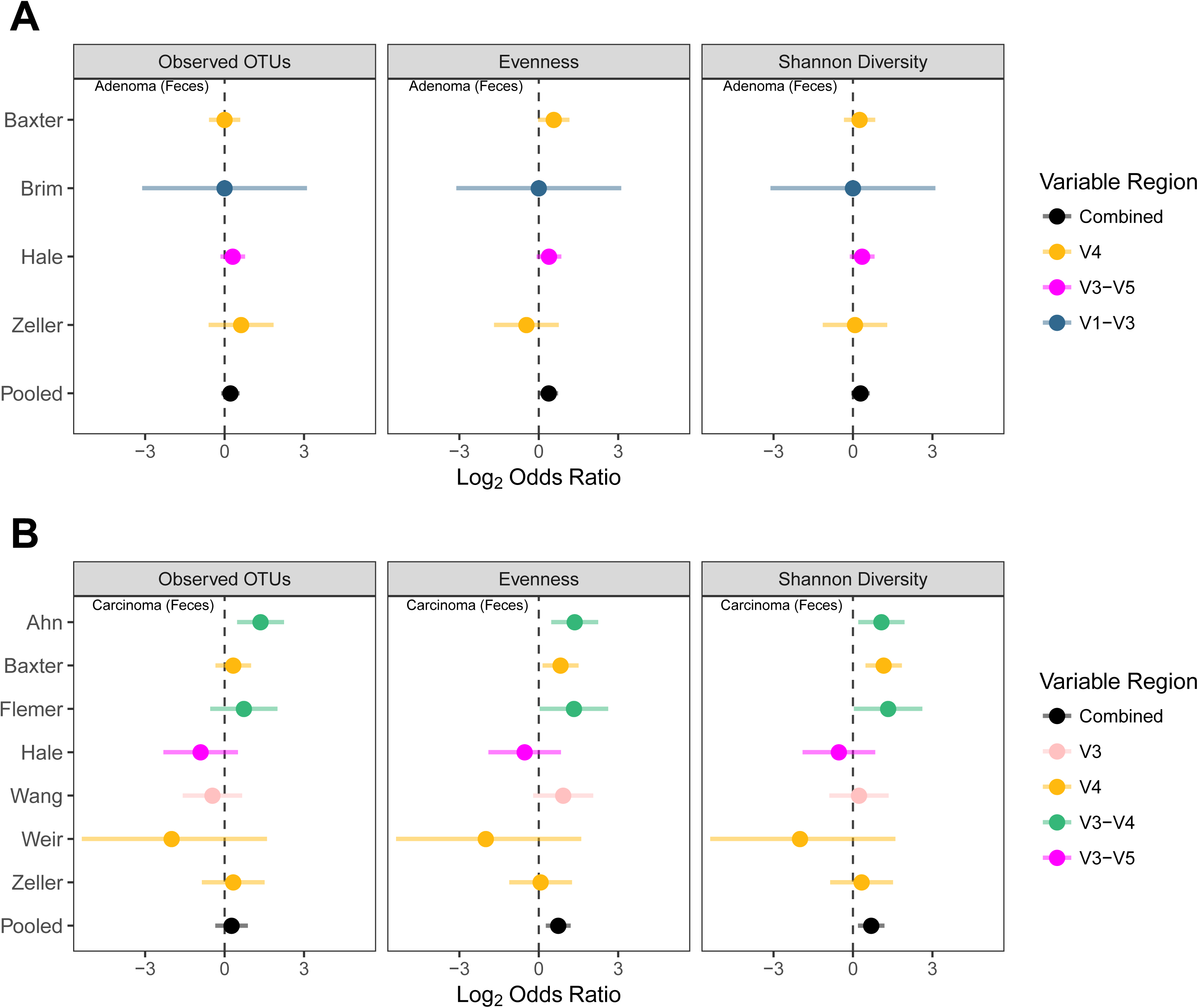
Comparison of odds ratios calculated using alpha diversity community metrics associated with the presence of adenomas (A) or carcinoma (B) relative to those in individuals with normal colons using data collected from stool samples.

Similar to our analysis of sequences obtained from fecal samples, we repeated the analysis using sequences obtained from colon tissue. There were no significant differences in richness, evenness, or diversity as disease severity progressed from control to adenoma to carcinoma (P-values > 0.05). We next analyzed the ORs, for matched (i.e. where unaffected tissue and tumors were obtained from the same individual) and unmatched (i.e. where unaffected tissue and tumor tissue were not obtained from the same individual) tissue samples. The ORs for adenoma and carcinoma were not significantly different from 1.0 for any measure (P-values > 0.05) [Figure S1 & Table S1]. This is likely due to the combination of a small effect size and the relatively small number of studies and size of studies used in the analysis.

### Disease progression is associated with changes in community structure

Based on the differences in evenness and diversity, we next asked whether there were community-wide differences in the structure of the communities associated with different disease stages. We identified significant bacterial community differences in the feces of patients with adenomas relative to those with normal colons in 1 of 4 studies and in patients with carcinomas relative to those with normal colons in 6 of 7 studies (PERMANOVA; P-value < 0.05) [Table S2]. Similar to the analyses using fecal samples, there were significant differences in the bacterial community structure of subjects with normal colons and those with adenomas (1 of 2 studies) and carcinomas (1 of 3 studies) [Table S2]. For studies that used matched samples, we did not observe any differences in bacterial community structures [Table S2]. Combined, these results indicate that there were consistent and significant community-wide changes in the fecal community structure of subjects with carcinomas. However, the signal observed in subjects with adenomas or when using tissue samples was not as consistent. This is likely due to a smaller effect size or the relatively small sample sizes among the studies that characterized the tissue microbiota.

### Individual taxa are associated with significant ORs for carcinomas

We next identified those taxa that had ORs that were significantly associated with having a normal colon or the presence of adenomas or carcinomas. No taxa had a significant OR for the presence of adenomas when we used data collected from fecal or tissue samples (Table S3 & S4). In contrast, 8 taxa had significant ORs for the presence of carcinomas using data from fecal samples. Of these, 4 are commonly associated with the oral cavity: *Fusobacterium* (OR=2.74 (1.95 - 3.85)), *Parvimonas* (OR=3.07 (2.11 - 4.46)), *Porphyromonas* (OR=3.2 (2.26 - 4.54)), and *Peptostreptococcus* (OR=7.11 (3.84 - 13.17)) [Table S3]. The other 4 were *Clostridium XI* (OR=0.65 (0.49 - 0.86)), *Enterobacteriaceae* (OR=1.79 (1.33 - 2.41)), *Escherichia* (OR=2.15 (1.57 - 2.95)), and *Ruminococcus* (OR=0.63 (0.48 - 0.83)). Among the data collected from tissue samples, only unmatched carcinoma samples had taxa with a significant OR. Those included *Dorea* (OR=0.35 (0.22 - 0.55)), *Blautia* (OR=0.47 (0.3 - 0.73)), and *Weissella* (OR=5.15 (2.02 -13.14)). Mouth-associated genera were not significantly associated with a higher OR for carcinoma in tissue samples [Table S4]. For example, *Fusobacterium* had an OR of 3.98 (1.19 - 13.24); however, due to the small number of studies and considerable variation in the data, the Benjimani-Hochberg corrected P-value was 0.93 [Table S4]. It is interesting to note that *Ruminococcus* and members of *Clostridium XI* in fecal samples and *Dorea* and *Blautia* in tissue had ORs that were significantly less than 1.0, which suggests that these populations are protective against the development of carcinomas. Overall, there was no overlap in the taxa with significant OR between fecal and tissue samples.

### Individual taxa with a significant OR do a poor job of differentiating subjects with normal colons and those with carcinoma

We next asked whether those taxa that had a significant OR associated with having a normal colon or carcinomas could be used individually, to classify subjects as having a normal colon or carcinomas. OR values were caluclated based on whether the relative abundance for a taxon in a subject was above or below the median relative abundance for that taxon across all subjects in a study. To measure the ability of these taxa to classify individuals we instead generated receiver operator characteristic (ROC) curves for each taxon in each study and calculated the area under the curve (AUC). This allowed us to use a more fluid relative abundance threshold for classifying individuals by their disease status. Using data from fecal samples, the 8 taxa did no better at classifying the subjects than one would expect by chance (i.e. AUC=0.50) [Figure 3A]. The taxa that performed the best included *Clostridium XI*, *Ruminococcus*, and *Escherichia*. However, these had median AUC values less than 0.588 indicating their limited value as biomarkers when used individually. Likewise, in unmatched tissue samples the 3 taxa with significant OR taxa had AUC values that were marginally better than one would expect by chance [Figure 3B]. The relative abundance of *Dorea* was the best predictor of carcinomas and its median AUC was only 0.62. These results suggest that although these taxa are associated with a significant OR for the presences of carcinomas, they do a poor job of classifying a subject’s disease status when used individually.

**Figure 3:**
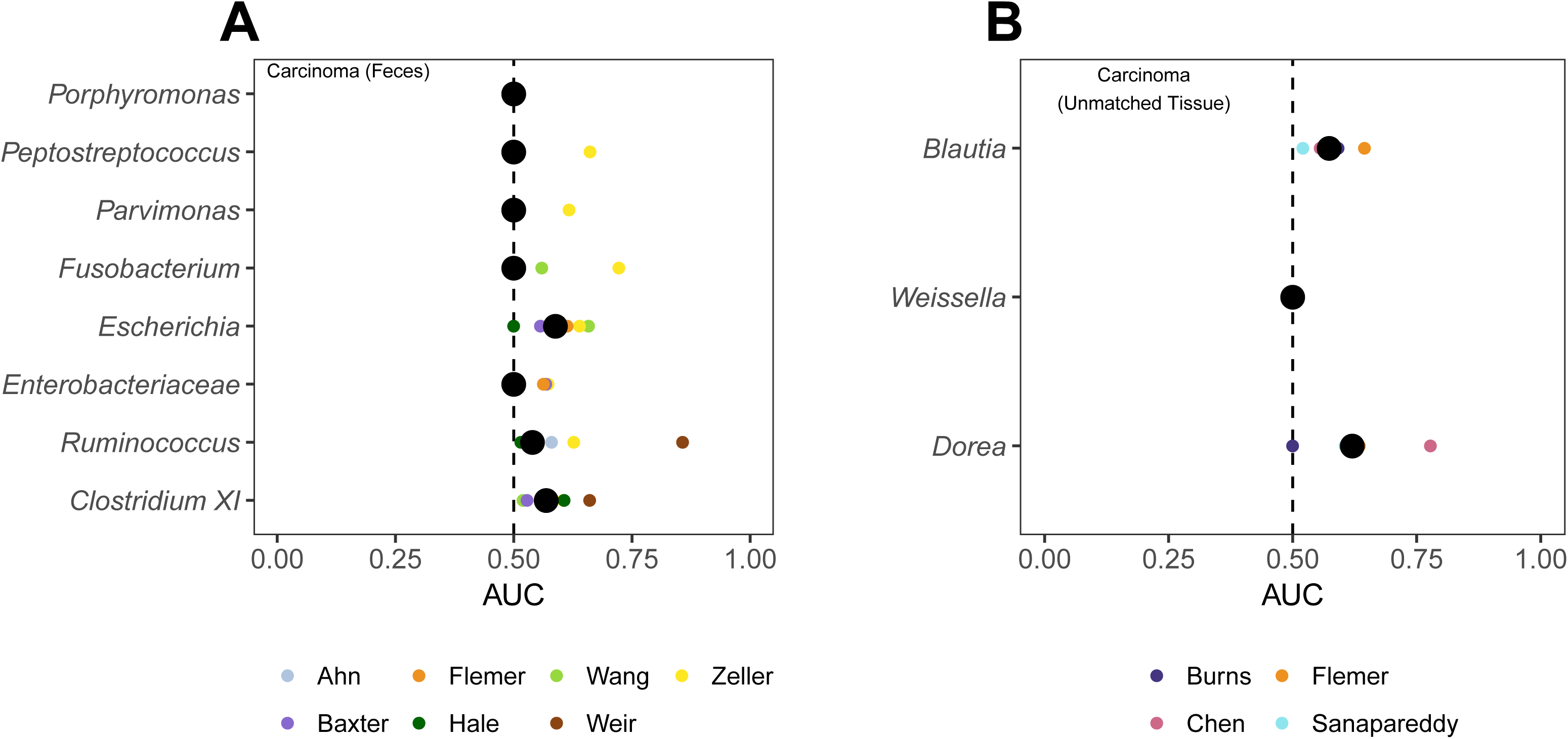
AUC values when classifing individuals as having normal colons or carcinomas using taxa with significant ORs when using stool samples (A) and unmatched tissue samples (B). We did not identify any taxa as having a significant OR to differentiate individuals with normal colons and adenomas or using matched tissue samples. The large black circles represent the median AUC of all studies and the smaller circles represent the individual AUC for a particular study. The dotted line denotes an AUC of 0.5.

### Combined taxa model classifies subjects better than using individual taxa

Instead of attempting to classify subjects based on individual taxa, next we combined information from the individual taxa and evaluated the ability to classify a subject’s disease status using Random Forest models. For data from fecal samples, the combined model had an AUC of 0.75, which was significantly higher than any of the AUC values for the individual taxa (P-value < 0.033). When this approach was used to train models using data from each study, the most important taxa were *Ruminococcus* and *Clostridium XI* [Figure 4A]. Similarly, using data from the unmatched tissue samples, the combined model had an AUC of 0.77, which was significantly higher than the AUC values for classifying based on the relative abundances of *Blautia* and *Weissella* individually (P-value < 0.037). Both *Dorea* and *Blautia* were the most important taxa in the tissue-based models [Figure 4B]. Pooling the information from the taxa with significant ORs resulted in models that outperformed classifications made using the same taxa individually.

**Figure 4:**
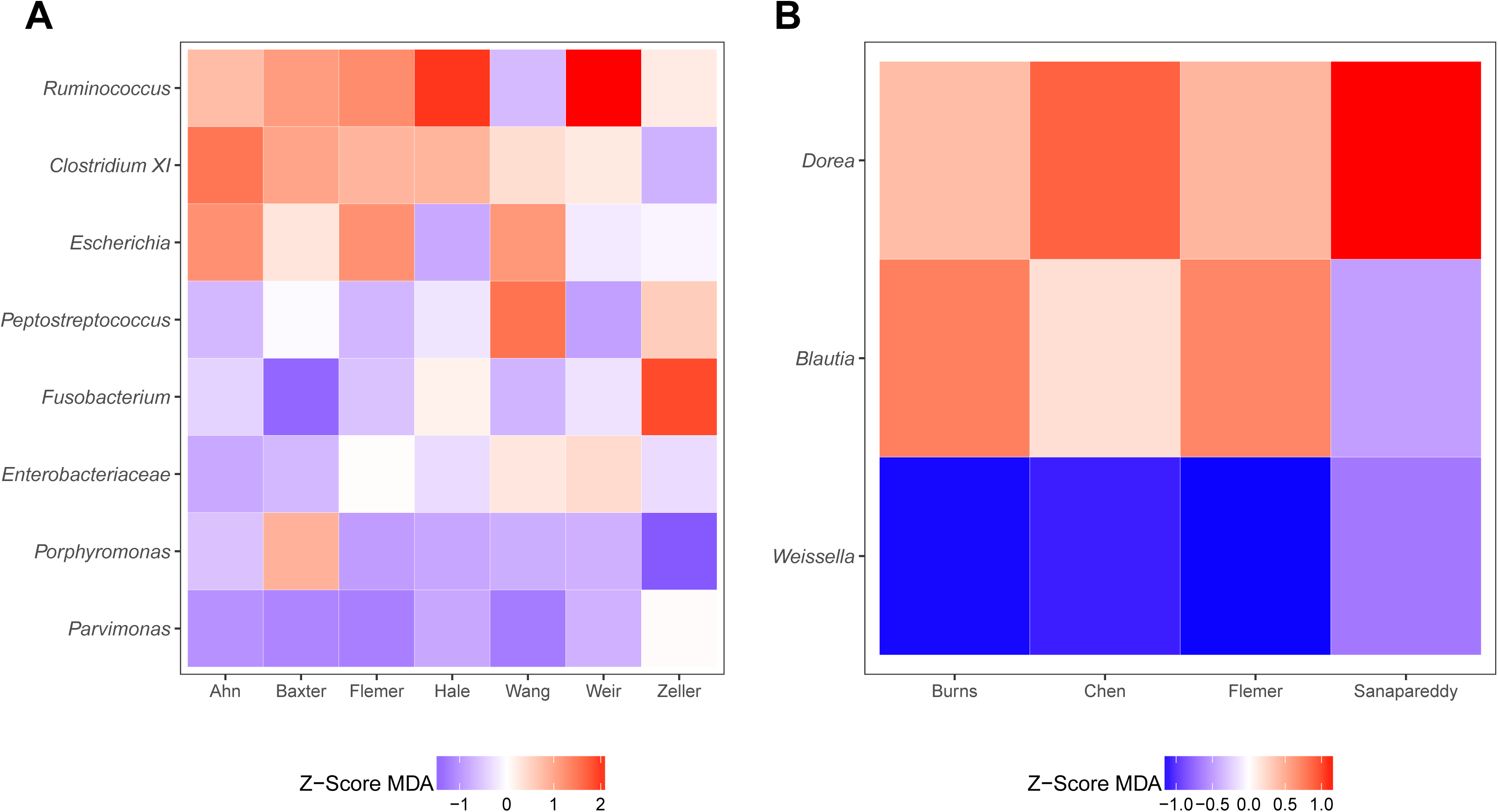
Relative importance of taxa with significant ORs in Random Forest models for differentiating between individuals with normal colons and carcinomas using stool samples (A) or unmatched tissue samples (B). The colors indicate the z-transformed (i.e. mean of 0.0 and standard deviation of 1.0) mean decrease in accuracy values calculated from the model for each study. The taxa are ranked by their mean z-score-transformed mean decrease in accuracy.

### Performance of models based on taxa relative abundance in full community is better than that of models based on taxa with significant ORs

Next, we asked whether a Random Forest classification model built using all of the taxa found in the communities would outperform the models generated using those taxa with a significant OR. Similar to our inability to identify taxa associated with a significant OR for the presence of adenomas, the median AUCs to classify subjects as having normal colons or having adenomas using data from fecal or tissue samples were only marginally better than 0.5 for any study (median AUC=0.549 (range: 0.367 - 0.971)) [Figure 5A & S2A]. In contrast, the models for classifying subjects as having normal colons or having carcinomas using data from fecal or tissue samples yielded AUC values meaningfully higher than 0.5 [Figure 5B & S2B-C]. When we compared the models based on all of the taxa in a community to models based on the taxa with significant ORs, the results were mixed. Using the data from fecal samples, we found that the AUC for 6 of 7 studies were an average of 14.8% higher and AUC for the Flemer study was 0.54% lower when using the relative abundance data from all taxa relative to using the relative abundance of only the taxa with significant ORs. The overall improvement in performance was statistically significant (mean=12.61%, one-tailed paired T-test; P-value=0.005). Among the models trained using data from fecal samples, *Bacteroides* and *Lachnospiraceae* were the most common taxa in the top 10% mean decrease in accuracy across studies [Figure S3]. Using data from unmatched tissue samples to train classification models, we found that the AUC of studies was an average 19.11% higher when we used all of the taxa rather than the 3 taxa with significant ORs (one-tailed paired T-test; P-value=0.03). For the models trained using data from unmatched tissue samples, *Lachnospiraceae*, *Bacteroidaceae*, and *Ruminococcaceae* were the most common taxa in the top 10% mean decrease in accuracy across studies [Figure S4]. Although the models trained using those taxa with a significant OR perform well for classifying individuals with and without carcinomas, models trained using data from the full community perform better.

**Figure 5:**
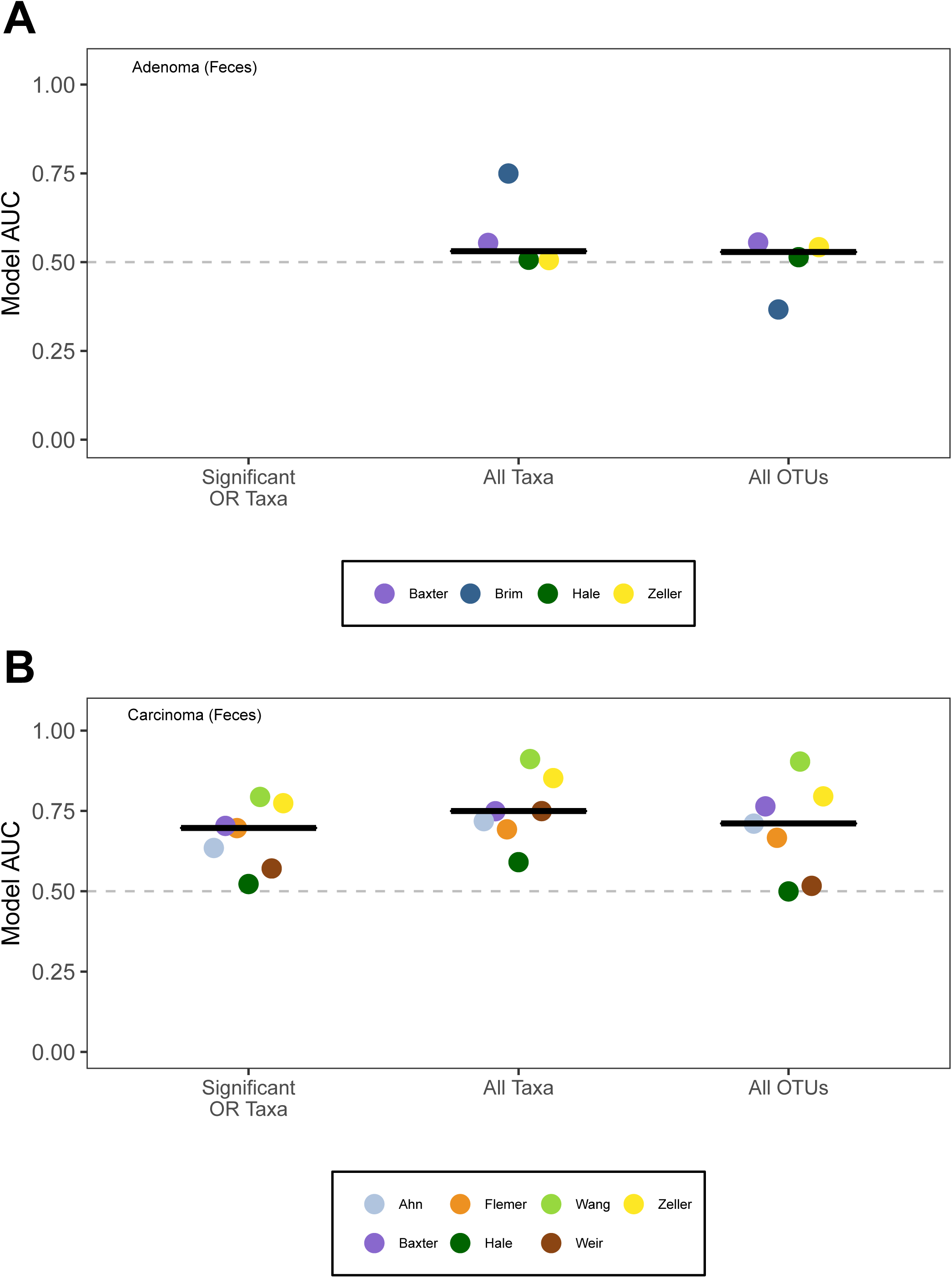
Comparison of Random Forest modeling approaches to classify individuals as having normal colons or adenomas (A) or carcinomas (B) when training the models using the taxa with significant ORs, all taxa in a community, or all OTUs in a community when using stool samples. No taxa had a significant OR associated with the presence of adenomas using stool samples. The black line represents the median AUC for the respective group. The dashed gray line indicates an AUC of 0.5.

### Performance of models based on OTU relative abundances are not significantly better than those based on taxa with significant ORs

The previous models were based on relative abundance data where sequences were classified to coarse taxonomic assignments (i.e. typically genus or family level). To determine whether model performance improved with finer scale classification, we assigned sequences to operational taxonomic units (OTUs) where the similarity among sequences within an OTU was more than 97%. We again found that classification models built using all of the sequence data for a community did a poor job of differentiating between subjects with normal colons and those with adenomas (median AUC: 0.53 (0.37- 0.56)). However, they did a good job of differentiating between subjects with normal colons and those with carcinomas (median AUC: 0.71 (0.50- 0.90)). The OTU-based models performed similarly to those constructed using the taxa with significant ORs (one-tailed paired T-test; P-value=0.979) and those using all taxa (one-tailed paired T-test; P-value=0.184) [Figure 4]. Among the OTUs that had the highest mean decrease in accuracy for the OTU-based models, we found that OTUs that affiliated with all of the 8 taxa that had a significant OR were within the top 10% for at least one study. This result was surprising as it indicated that a finer scale classification of the sequences and thus a larger number of features to select from, did not yield improved classification of the subjects.

### Generalizability of taxon-based models trained on one dataset to the other datasets

Considering the good performance of the Random Forest models trained using the relative abundance of taxa with significant ORs and models trained using the relative abundance of all taxa, we next asked how well the models would perform when given data from a different cohort. For instance, if a model was trained using data from the Ahn study, we wanted to know how well it would perform using the data from the Baxter study. The models trained using the taxa with significant ORs all had a higher median AUC than the models trained using all of the taxa when tested on the other datasets [Figure 6 & S5]. As might be expected, the difference between the performance of the modeling approaches appeared to vary with the size of the training cohort (R^2^=0.66) [Figure 6]. These data suggest that given a sufficient number of subjects with normal colons and carcinomas, Random Forest models trained using a small number of taxa can accurately classify individuals from a different cohort.

**Figure 6:**
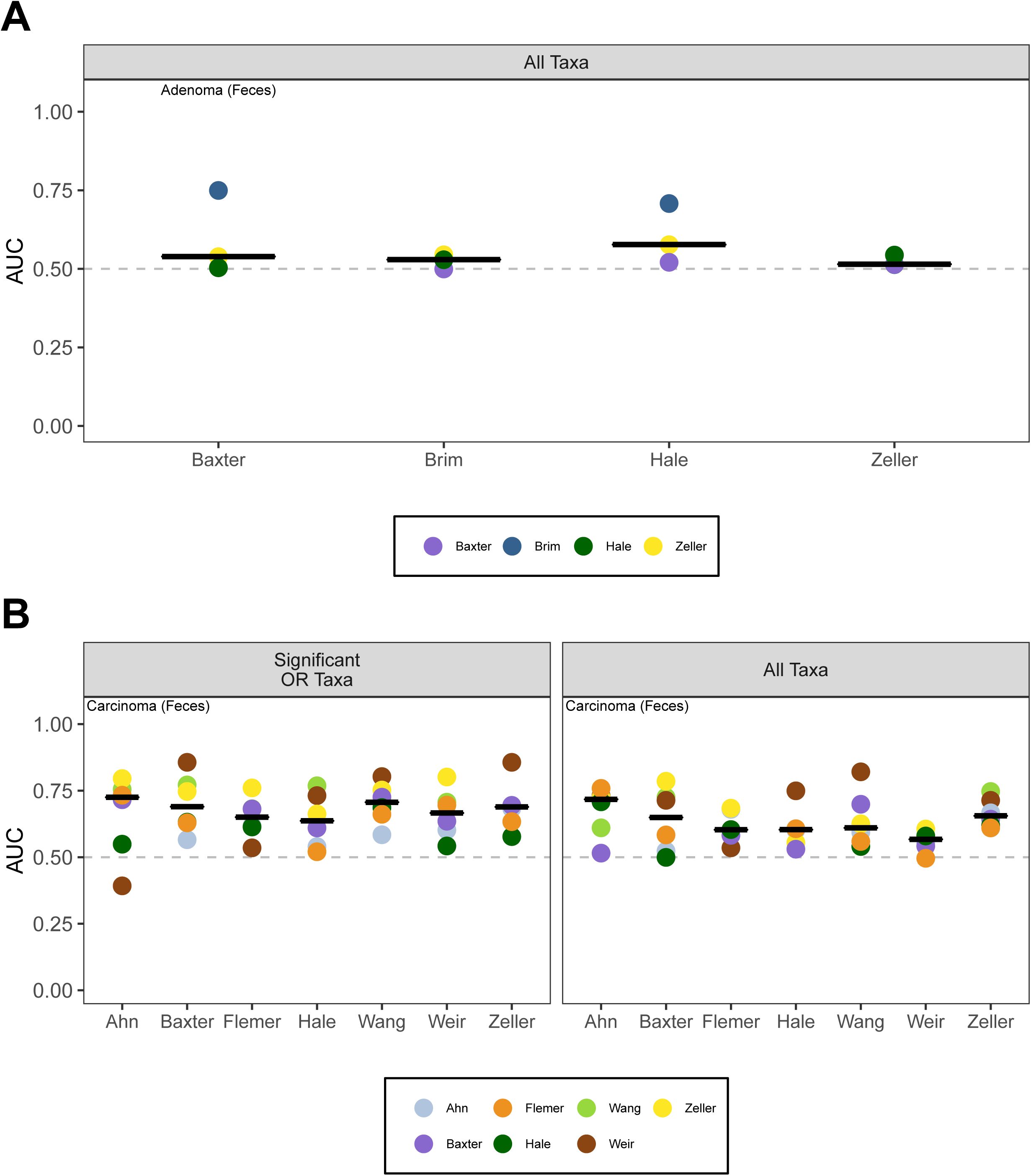
Testing of Random Forest models to classify individuals as having normal colons or adenomas (A) or carcinomas (B) when using sequence data obtained from stool samples. Models were trained on data from each study (Figure 5) and tested on the other studies. The black lines represent the median AUC of all test AUCs for a specific study. The dashed gray line represents the AUC at 0.5.

## Discussion

We performed a meta-analysis to identify and validate microbiota-based biomarkers that could be used to classify individuals as having normal colons or colonic tumors using fecal or tissue samples. To our surprise, Random Forest classification models constructed to differentiate individuals with normal colons from those with carcinomas using a subset of the community performed well relative to models constructed using the full communities. When we applied the models trained on each dataset to the other datasets in our study, we found that the models trained using the subset of the communities performed better than those using the full communities. These models were trained using data in which sequences were assigned to bacterial taxa using a classifier that typically assigned sequences to the family or genus level. When we attempted to improve the specificity of the classification by using an OTU-based approach the resulting models performed as well as those constructed using coarse taxonomic assignments. These results are significant because they strengthen the growing literature indicating a role for the colonic microbiota in tumorigenesis, as a potential tool as a non-invasive diagnostic, and for assessing risk of disease and recurrence (9, 12, 30).

Fine scale classification of sequences into OTUs did not improve our classification models. This was also tested in earlier efforts to use shotgun metagenomic data to classify individuals as having normal colons or tumors; however, it was shown that analyses performed using shotgun metagenomic data did not perform better than using 16S rRNA gene sequencing data (31). We hypothesize that fine scale classification may not result in better classification because distribution of microbiota between individuals is patchy. In contrast, models using coarser taxonomic assignments will pool the fine scale diversity, resulting in less patchiness and better classification. Furthermore, the ability of models trained using a subset of the community to outperform those using the full community when testing the models on the other datasets may also be a product of the patchiness of the human-associated microbiota. The models based on the 8 taxa that had significant ORs used taxa that were found in every study and tended to have higher relative abundances. Similar to the OTU-based models, those models based on the full community taxonomy assignments were still sensitive to the patchy distribution of taxa. Regardless, it is encouraging that a collection of 8 taxa could reliably classify individuals as having carcinomas considering the differences in cohorts, DNA extraction procedures, regions of the 16S rRNA gene, and sequencing methods.

When used to classify individuals with carcinomas, the taxa with significant ORs could not reliably classify individuals on their own [Figure 3]. This result further supports the hypothesis that carcinoma-associated microbiota have a patchy distribution. Two individuals may have had the same classification, based on the relative abundance of different populations within this group of 8 taxa. Although these results only reflect associations with disease, it is tempting to hypothesize that the patchiness is indicative of distinct mechanisms of exacerbating tumorigenesis or that multiple taxa have the same mechanism of exacerbating tumorigenesis. For example, strains of *Escherichia coli* and *Fusobacterium nucleatum* have been shown to worsen inflammation in mouse models of tumorigenesis (5, 6, 21). In contrast to the patchiness of the taxa that were positively associated with carcinomas, potentially beneficial taxa had a more consistent association [Figure 6]. This result was particularly interesting because members of these taxa (i.e. *Ruminococcus* and *Clostridium XI* in fecal samples and *Dorea* and *Blautia* in tissue) are thought to be beneficial due to their involvement in production of anti-inflammatory short chain fatty acids (32–34).

All of the adenoma classification models performed poorly, which is consistent with previous studies (27, 30). However, the classification results are at odds with results of the multitarget microbiota test (MMT) from Baxter, et al. (12) who observed an AUC of 0.755 when the test was applied to individuals with adenomas. There are two major differences between the models generated in this meta-analysis and that analysis. The MMT attempted to classify individuals as having a normal colon or having colonic lesions (i.e. adenomas or carcinomas) and not adenomas alone. Further, the MMT incorporated fecal immunoglobulin test (FIT) data while our models only used 16S rRNA gene sequencing data. Because FIT data were not available for the other studies in our meta-analysis, it was not possible to validate the MMT approach. The ability to differentiate between individuals with and without adenomas is an important problem since early detection of tumors is critical to patient survivorship. However, it is possible that we might have been able to detect differences in the bacterial community if individuals with non-advanced and advanced adenomas were separated. This is a clinically relevant distinction since advanced adenomas are at highest risk of progressing to carcinomas. The initial changes of the microbiota during tumorigenesis could be focal to where the initial adenoma develops and would not be easily assessed using fecal samples from an individual with non-advanced adenomas. Unfortunately, distinguishing between individuals with advanced and non-advanced adenomas was not possible in our meta-analysis since the studies did not provide the clinical data needed to make that distinction.

Fecal samples represent a non-invasive approach to assess the structure of the gut microbiota and are potentially useful for diagnosing individuals as having colonic tumors. However, they do not reflect the structure of the mucosal microbiota (35). Regardless, the taxa that were the most important in the feces-based models overlapped with those from the models trained using the data from unmatched and matched colon tissue samples [Figure S3]. Mucosal biopsies are preferred for focused mechanistic studies and have offered researchers the opportunity to sample healthy and diseased tissue from the same individuals (i.e. matched) using each individual as their own control or in a cross-sectional design (i.e. unmatched). Because obtaining these samples is invasive, carries risks to the individual, and is expensive, studies investigating the structure of the mucosal microbiota generally have a limited number of participants. Thus, it was not surprising that tissue-based studies did not provide clearer associations between the mucosal microbiota and the presence of tumors. Interestingly, *Fusobacterium*, which has received increased attention for its potential role in tumorigenesis (6) was not consistently identified across the studies in our meta-analysis which is consistent with a recent replicability study (36). This could be due to the relatively small number of individuals in the limited number of studies. The classification models trained using the tissue-based data performed well when tested with the training data (Figure S4), but performed poorly when tested on the other tissue-associated datasets (Figure S5). Disturbingly, taxa that are commonly associated with reagent contamination (e.g. *Novosphingobium*, *Acidobacteria Gp2*, *Sphingomonas*, etc.) were detected within the tissue datasets. Such contamination is common in studies where there is relatively low bacterial biomass (37). The lack of replication among the tissue-based biomarkers may be a product of the relatively small number of studies and individuals per study and possible reagent contamination.

Among the fecal sample data, we failed to identify several notable populations that are commonly associated with carcinomas including an enterotoxigenic strain of *Bacteroides fragilis* (ETBF) and *Streptococcus gallolyticus* subsp. *gallolyticus* (22, 24). ETBF have been found in tumors in the proximal colon where they tend to form biofilms (20, 38). Considering DNA from bacteria that are more prevalent in the proximal colon may be degraded by the time it leaves the body, it is not surprising that we failed to identify a significant OR for *Bacteroides* with carcinomas. In addition, since our approach could only classify sequences to the genus level and there are likely multiple *Bacteroides* populations in the colon, it is possible that sequences from ETBF and non-oncogenic *Bacteroides* were pooled. This would then reduce the OR between *Bacterioides* and whether an individual had carcinomas. It is also necessary to distinguish between populations that are biomarkers for a disease and those that are known to cause disease. Although the latter have been shown to have a causative role, they may appear at low relative abundance, be found in specific locations, or may have a highly patchy distribution among affected individuals.

Meta-analyses are a useful tool in microbiome research because they can demonstrate whether a result can be replicated and facilitate new discoveries by pooling multiple independent investigations. There have been several meta-analyses similar to this study that have sought biomarkers for obesity (39–41), inflammatory bowel disease (40), and colorectal cancer (28). Considering microbiome research is particularly prone to hype and overgeneralization of results (42), these analyses are critical. Meta-analyses are difficult to perform because the underlying 16S rRNA gene sequence data are not publicly available, metadata are missing, incomplete, or vague, sequence data are of poor quality or derived by non-standard approaches, and the original studies may be significantly underpowered. Reluctance to publish negative results (i.e. the “file drawer effect”) is also likely to skew our understanding of the relationship between microbiota and disease. Better attention to these specific issues will increase the reproducibility and replicability of microbiota studies and make it easier to perform these crucial meta-analyses. Moving forward, meta-analyses will be important tools to help aggregate and find commonalities across studies when investigating the microbiota in the context of a specific disease (28, 39–41).

Our meta-analysis suggests a strong association between the gut microbiota and colon tumorigenesis. By aggregating the results from studies that sequenced the 16S rRNA gene from fecal and tissue samples, we are able to provide evidence supporting the use of microbial biomarkers to diagnose the presence of colonic tumors. Further development of microbial biomarkers should focus on including other biomarkers (e.g. FIT), better categorizing of people with adenomas, and expanding datasets to include larger numbers of individuals. Based on prior research into the physiology of the biomarkers we identified, it is likely that they have a causative role in tumorigenesis. Their patchy distribution across individuals suggests that there are either multiple mechanisms causing disease or a single mechanism (e.g. inflammation) that can be mediated by multiple, diverse bacteria.

## Methods

### Datasets

The studies used for this meta-analysis were identified through the review articles written by Keku, et al. (43) and Vogtmann, et al. (44). Additional studies, not mentioned in those reviews were obtained based on the authors’ knowledge of the literature. Studies were included that used tissue or feces as their sample source for 454 or Illumina 16S rRNA gene sequencing. A significant number of studies (N=12) were excluded from the meta-analysis because they did not have publicly available sequences, did not use 454 or Illumina sequencing platforms, or did not have metadata that the authors were able to share. We were able to obtain sequence data and metadata from the following studies: Ahn, et al. (11), Baxter, et al. (12), Brim, et al. (29), Burns, et al. (15), Chen, et al. (13), Dejea, et al. (20), Flemer, et al. (17), Geng, et al. (19), Hale, et al. (27), Kostic, et al. (45), Lu, et al. (26), Sanapareddy, et al. (25), Wang, et al. (14), Weir, et al. (23), and Zeller, et al. (16). The Zackular (46) study was excluded because the individuals studied were included within the larger Baxter study (12). The Kostic study was excluded because after we processed the sequences, all of the case samples had 100 or fewer sequences. The final analysis included 14 studies (Tables 1 and 2). There were seven studies with only fecal samples (Ahn, Baxter, Brim, Hale, Wang, Weir, and Zeller), five studies with only tissue samples (Burns, Dejea, Geng, Lu, Sanapareddy), and two studies with both fecal and tissue samples (Chen and Flemer). After curating the sequences, 1737 fecal samples and 492 tissue samples remained in the analysis [Tables 1 and 2].

### Sequence Processing

Raw sequence data and metadata were primarily obtained from the Sequence Read Archive (SRA) and dbGaP. Other sequence and metadata were obtained directly from the authors (n=4, (17, 23, 25, 27)). Each dataset was processed separately using mothur (v1.39.3) using the default quality filtering methods for both 454 and Illumina sequence data (47). If it was not possible to use the defaults because the trimmed sequences were too short, then the stated quality cut-offs from the original study were used. Chimeric sequences were identified and removed using VSEARCH (48). The curated sequences were assigned to OTUs at 97% similarity using the OptiClust algorithm (49) and classified to the deepest taxonomic level that had 80% support using the naïve Bayesian classifier trained on the RDP taxonomy outline (version 14, (50)).

### Community analysis

We calculated alpha diversity metrics (i.e. OTU richness, evenness, and Shannon diversity) for each sample. Within each dataset, we ensured that the data followed a normal distribution using power transformations. Using the transformed data, we tested the hypothesis that individuals with normal colons, adenomas, and carcinomas had significantly different alpha diversity metrics using linear mixed-effect models. We also calculated the OR for each study and metric by considering any value above the median alpha diversity value to be positive. We measured the dissimilarity between individuals by calculating the pairwise Bray-Curtis index and used PERMANOVA (51) to test whether individuals with normal colons were significantly different from those with adenomas or carcinomas. Finally, after binning sequences into the deepest taxa that the naïve Bayesian classifier could calssify the sequences, we quantified the ORs for individuals having an adenoma or carcinoma and corrected for multiple comparisons using the Benjamini-Hochberg method (52). Again, for each taxon, if the relative abundance was greater than the median relative abundance for that taxon in the study, the individual was considered to be positive.

### Random Forest classification analysis

To classify individuals as having normal colons or tumors, we built Random Forest classification models for each dataset and comparison using taxa with significant ORs (after multiple comparison correction), all taxa, or OTUs. Because no taxa were identified as having a significant OR associated with adenomas using stool or tissue samples, classification models based on OR data were not constructed to classify individuals as having normal colons or adenomas. For all models, the value of trees included (i.e. ntree) was set to 500 and the number of variables that were randomly tested (i.e. mtry) was set to the square root of the number of taxa or OTUs within the model. Using the square root of the total number of features as the number of features to test has been found to reliably approximate the optimum value after model tuning (53). All fecal models were built using a 10-fold cross validation (CV) while tissue models were built using 5-fold CV due to study sample size. One exception to this were the models constructed using data from the Weir study, which was built using a 2-fold CV due to the small number of samples. For models constructed based on the taxa that had a significant OR or using all of the taxa, we trained the models using a single study and then tested on the remaining studies with AUCs recorded during both train and testing phases. For the models constructed using OTU data, 100 10-fold CVs were run to generate a range of AUCs that could be reasonably expected to occur. The average AUC from these 100 repeats was reported. The Mean Decrease in Accuracy (MDA), a measure of the importance of each taxon to the overall model, was used to rank the taxa used in each model.

### Statistical Analysis

All statistical analysis after sequence processing utilized the R (v3.4.3) software package (54). For OTU richness, evenness, and Shannon diversity analysis, values were power transformed using the rcompanion (v1.11.1) package (55) and Z-score normalized using the car (v2.1.6) package (56). Testing for OTU richness, evenness, and Shannon diversity differences utilized linear mixed-effect models to correct for study, repeat sampling of individuals (tissue only), and 16S rRNA gene sequence region used using the lme4 (v1.1.15) package (57). ORs were analyzed using both the epiR (v0.9.93) and metafor (v2.0.0) packages (58, 59) by assessing how many individuals with and without disease were above and below the overall median value within each specific study. OR significance testing utilized the chi-squared test. Community structure differences were calculated using the Bray-Curtis dissimilarity index and PERMANOVA was used to test for tumor-associated differences in structure with the vegan (v2.4.5) package (60). Random Forest models were built using both the caret (v6.0.78) and randomForest (v4.6.12) packages (61, 62). All figures were created using both ggplot2 (v2.2.1) and gridExtra (v2.3) packages (63, 64).

### Reproducible Methods

The analysis code can be found at https://github.com/ SchlossLab/Sze_CRCMetaAnalysis_mBio_2018. Unless otherwise mentioned, the accession number of raw sequences from the studies used in this analysis can be found directly in the respective batch file in the GitHub repository or in the original manuscript.

## Acknowledgements

The authors would like to thank all the study participants who were a part of each of the individual studies analyzed. We would also like to thank each of the study authors for making their sequencing reads and metadata available for use. Finally, we would like to thank the members of the Schloss lab for their valuable feedback and proofreading during the formulation of this manuscript.

---

**Figure S1: Comparison of Odds Ratios associated with normal colons or adenomas (A) or carcinomas (B) calculated using alpha diversity indices with sequence data generated from tissue samples.** The pooled results are from the aggregation of data across all studies. The horizontal lines indicate the 95% confidence interval for the OR.

**Figure S2: Comparison of Random Forest modeling approaches to classify individuals as having normal colons or adenomas (A) or carcinomas (B) when training the models using the taxa with significant ORs, all taxa in a community, or all OTUs in a community when using data from tissue samples.** No taxa had a significant OR associated with the presence of adenomas using tissue samples. The black line represents the median AUC for the respective group. The dashed gray line indicates an AUC of 0.5.

**Figure S3: Relative importance of taxa (A) and OTUs (B) in Random Forest models for differentiating between individuals with normal colons and carcinomas using stool samples.** These taxa and OTUs were among the top 10% most important features in each model. The colors indicate the z-transformed (i.e. mean of 0.0 and standard deviation of 1.0) mean decrease in accuracy values calculated from the model for each study. The taxa are ranked by their mean z-score-transformed mean decrease in accuracy.

**Figure S4: Relative importance of taxa (A, B) and OTUs (C, D) in Random Forest models for differentiating between individuals with normal colons and carcinomas using matched (A, C) and unmatched (B, D) tissue samples.** hese taxa and OTUs were among the top 10% most important features in each model. The colors indicate the z-transformed (i.e. mean of 0.0 and standard deviation of 1.0) mean decrease in accuracy values calculated from the model for each study. The taxa are ranked by their mean z-score-transformed mean decrease in accuracy.

**Figure S5: Testing of Random Forest models to classify individuals as having normal colons or adenomas (A) or carcinomas (B, C) when using sequence data obtained from tissue samples.** Models were trained on data from each study (Figure S5) and tested on the other studies. The black lines represent the median AUC of all test AUCs for a specific study. The dashed gray line represents the AUC at 0.5.

