## Supplementary Materials for "Leveraging Existing 16S rRNA Gene Surveys to Identify Reproducible Biomarkers in Individuals with Colorectal Tumors"

Supplemental

**Table S1: Comparison of odds ratios calculated using alpha diversity community metrics associated with the presence of adenomas or carcinoma relative to those in individuals with normal colons using data collected from tissue samples.**

| Odds Ratio | 95% CI (Lower Bound) | 95% CI (Upper Bound) | P-value | Measure | Tumor | Tissue Group |
|---|---|---|---|---|---|---|
| 1.82 | 0.95 | 3.49 | 0.07 | OTU Richness | Adenoma | Combined |
| 3.25 | 0.55 | 19.25 | 0.19 | Shannon Diversity | Adenoma | Combined |
| 3.47 | 0.68 | 17.70 | 0.13 | Evenness | Adenoma | Combined |
| 1.53 | 0.77 | 3.04 | 0.22 | OTU Richness | Carcinoma | Unmatched |
| 1.43 | 0.67 | 3.05 | 0.35 | Shannon Diversity | Carcinoma | Unmatched |
| 1.84 | 0.54 | 6.24 | 0.33 | Evenness | Carcinoma | Unmatched |
| 0.43 | 0.07 | 2.45 | 0.34 | OTU Richness | Carcinoma | Matched |
| 0.45 | 0.14 | 1.43 | 0.17 | Shannon Diversity | Carcinoma | Matched |
| 0.45 | 0.14 | 1.43 | 0.17 | Evenness | Carcinoma | Matched |

**Table S2: Comparison of community dissimilarity between individuals with normal colons and those with adenomas and carcinomas as calculated using Bray-Curtis distance and tested using PERMANOVA.**

| Study | Tumor | Sample Type | R2 | P-value |
|---|---|---|---|---|
| Brim | Adenoma | Feces | 0.059 | 0.7562 |
| Zeller | Adenoma | Feces | 0.021 | 0.0096 |
| Baxter | Adenoma | Feces | 0.003 | 0.3788 |
| Hale | Adenoma | Feces | 0.001 | 0.3658 |
| Wang | Carcinoma | Feces | 0.034 | 0.0001 |
| Weir | Carcinoma | Feces | 0.107 | 0.3431 |
| Ahn | Carcinoma | Feces | 0.010 | 0.0033 |
| Zeller | Carcinoma | Feces | 0.028 | 0.0003 |
| Baxter | Carcinoma | Feces | 0.007 | 0.0024 |
| Hale | Carcinoma | Feces | 0.002 | 0.7163 |
| Flemer | Carcinoma | Feces | 0.016 | 0.0460 |
| Lu | Adenoma | Tissue | 0.144 | 0.0001 |
| Flemer | Adenoma | Tissue | 0.018 | 0.0001 |
| Lu (Matched) | Adenoma | Tissue | 0.569 | 0.1000 |
| Sanapareddy | Carcinoma | Tissue | 0.025 | 0.0069 |
| Burns | Carcinoma | Tissue | 0.051 | 0.0995 |
| Flemer | Carcinoma | Tissue | 0.029 | 0.0001 |
| Chen | Carcinoma | Tissue | 0.064 | 0.2691 |
| Dejea (Matched) | Carcinoma | Tissue | 0.048 | 0.2515 |
| Geng (Matched) | Carcinoma | Tissue | 0.030 | 0.9816 |
| Burns (Matched) | Carcinoma | Tissue | 0.168 | 1.0000 |

**Table S3: ORs for individual taxa associated with individuals who had a normal colon or adenomas or carcinomas using data collected from stool.** The listed P-values were less than 0.05 prior to using a Benjimani-Hochberg correction for multiple comparisons.

| Taxon | Tumor | OR | 95% CI (Lower Bound) | 95% CI (Upper Bound) | P-value | BH |
|---|---|---|---|---|---|---|
| Clostridium_XlVb | Adenoma | 1.46 | 1.14 | 1.86 | 2.29e-03 | 2.20e-01 |
| Porphyromonas | Adenoma | 1.77 | 1.19 | 2.62 | 4.48e-03 | 2.20e-01 |
| Lachnospiraceae | Adenoma | 0.71 | 0.56 | 0.91 | 6.40e-03 | 2.20e-01 |
| Novosphingobium | Adenoma | 3.33 | 1.27 | 8.72 | 1.41e-02 | 2.92e-01 |
| Bacteroidales | Adenoma | 1.35 | 1.06 | 1.72 | 1.63e-02 | 2.92e-01 |
| Clostridium_XI | Adenoma | 0.75 | 0.59 | 0.95 | 1.92e-02 | 2.92e-01 |
| Clostridiaceae_1 | Adenoma | 0.71 | 0.54 | 0.95 | 1.99e-02 | 2.92e-01 |
| Lactococcus | Adenoma | 0.68 | 0.47 | 0.97 | 3.56e-02 | 4.59e-01 |
| Porphyromonas | Carcinoma | 3.20 | 2.26 | 4.54 | 6.73e-11 | 5.59e-09 |
| Peptostreptococcus | Carcinoma | 7.11 | 3.84 | 13.17 | 4.60e-10 | 1.91e-08 |
| Parvimonas | Carcinoma | 3.07 | 2.11 | 4.46 | 3.80e-09 | 1.05e-07 |
| Fusobacterium | Carcinoma | 2.74 | 1.95 | 3.85 | 5.54e-09 | 1.15e-07 |
| Escherichia.Shigella | Carcinoma | 2.15 | 1.57 | 2.95 | 2.20e-06 | 3.65e-05 |
| Enterobacteriaceae | Carcinoma | 1.79 | 1.33 | 2.41 | 1.30e-04 | 1.80e-03 |
| Ruminococcus | Carcinoma | 0.63 | 0.48 | 0.83 | 1.19e-03 | 1.41e-02 |
| Clostridium_XI | Carcinoma | 0.65 | 0.49 | 0.86 | 2.94e-03 | 3.05e-02 |
| Roseburia | Carcinoma | 0.60 | 0.41 | 0.88 | 8.98e-03 | 8.28e-02 |
| Clostridium_XlVb | Carcinoma | 1.45 | 1.09 | 1.94 | 1.17e-02 | 9.72e-02 |
| Clostridiaceae_1 | Carcinoma | 0.67 | 0.48 | 0.93 | 1.53e-02 | 1.15e-01 |
| Campylobacter | Carcinoma | 1.76 | 1.10 | 2.82 | 1.88e-02 | 1.30e-01 |
| Anaerococcus | Carcinoma | 2.50 | 1.14 | 5.47 | 2.22e-02 | 1.42e-01 |
| Desulfovibrio | Carcinoma | 1.45 | 1.05 | 2.00 | 2.54e-02 | 1.50e-01 |
| Veillonellaceae | Carcinoma | 1.53 | 1.04 | 2.26 | 3.28e-02 | 1.79e-01 |
| Lachnospiraceae | Carcinoma | 0.69 | 0.49 | 0.97 | 3.44e-02 | 1.79e-01 |

**Table S4: ORs for individual taxa associated with individuals who had a normal colon or adenomas or carcinomas using data collected from tissue samples.** The listed P-values were less than 0.05 prior to using a Benjimani-Hochberg correction for multiple comparisons.

| Taxon | Tumor | Tissue Group | OR | 95% CI (Lower Bound) | 95% CI (Upper Bound) | P-value | BH |
|---|---|---|---|---|---|---|---|
| Lachnospiraceae | Adenoma | Combined | 0.27 | 0.14 | 0.56 | 3.23e-04 | 7.17e-02 |
| Pseudomonas | Adenoma | Combined | 3.73 | 1.73 | 8.05 | 8.13e-04 | 7.17e-02 |
| Howardella | Adenoma | Combined | 3.69 | 1.72 | 7.95 | 8.30e-04 | 7.17e-02 |
| Rothia | Adenoma | Combined | 2.87 | 1.48 | 5.58 | 1.82e-03 | 1.18e-01 |
| Blautia | Adenoma | Combined | 0.39 | 0.20 | 0.77 | 6.96e-03 | 3.61e-01 |
| Enterobacter | Adenoma | Combined | 3.42 | 1.28 | 9.12 | 1.42e-02 | 4.65e-01 |
| Puniceicoccaceae | Adenoma | Combined | 2.94 | 1.22 | 7.10 | 1.61e-02 | 4.65e-01 |
| Erysipelotrichaceae | Adenoma | Combined | 2.27 | 1.16 | 4.42 | 1.62e-02 | 4.65e-01 |
| Streptococcus | Adenoma | Combined | 2.27 | 1.16 | 4.42 | 1.62e-02 | 4.65e-01 |
| Lactococcus | Adenoma | Combined | 3.24 | 1.17 | 9.00 | 2.37e-02 | 6.15e-01 |
| Micrococcaceae | Adenoma | Combined | 3.91 | 1.17 | 13.06 | 2.65e-02 | 6.24e-01 |
| Shewanella | Adenoma | Combined | 2.05 | 1.05 | 3.98 | 3.46e-02 | 6.66e-01 |
| Phascolarctobacterium | Adenoma | Combined | 2.03 | 1.05 | 3.93 | 3.57e-02 | 6.66e-01 |
| Achromobacter | Adenoma | Combined | 7.10 | 1.14 | 44.44 | 3.61e-02 | 6.66e-01 |
| Anaerostipes | Adenoma | Combined | 0.10 | 0.01 | 0.89 | 3.96e-02 | 6.66e-01 |
| Neisseria | Adenoma | Combined | 2.09 | 1.00 | 4.35 | 4.88e-02 | 6.66e-01 |
| Fusobacterium | Adenoma | Combined | 2.85 | 1.00 | 8.12 | 4.94e-02 | 6.66e-01 |
| Dorea | Carcinoma | Unmatched | 0.35 | 0.22 | 0.55 | 3.96e-06 | 4.40e-04 |
| Weissella | Carcinoma | Unmatched | 5.15 | 2.02 | 13.14 | 5.96e-04 | 2.88e-02 |
| Blautia | Carcinoma | Unmatched | 0.47 | 0.30 | 0.73 | 7.79e-04 | 2.88e-02 |
| Campylobacter | Carcinoma | Unmatched | 2.13 | 1.23 | 3.67 | 6.57e-03 | 1.80e-01 |
| Leptotrichia | Carcinoma | Unmatched | 2.71 | 1.30 | 5.67 | 8.10e-03 | 1.80e-01 |
| Parvimonas | Carcinoma | Unmatched | 1.94 | 1.17 | 3.21 | 1.05e-02 | 1.95e-01 |
| Clostridiaceae_1 | Carcinoma | Unmatched | 1.99 | 1.15 | 3.45 | 1.40e-02 | 2.15e-01 |
| Flavobacteriaceae | Carcinoma | Unmatched | 2.34 | 1.17 | 4.69 | 1.64e-02 | 2.15e-01 |
| Ruminococcus2 | Carcinoma | Unmatched | 0.28 | 0.10 | 0.81 | 1.81e-02 | 2.15e-01 |
| Clostridium_XlVb | Carcinoma | Unmatched | 0.60 | 0.39 | 0.92 | 1.95e-02 | 2.15e-01 |
| Corynebacterium | Carcinoma | Unmatched | 0.52 | 0.29 | 0.92 | 2.41e-02 | 2.15e-01 |
| Finegoldia | Carcinoma | Unmatched | 0.43 | 0.21 | 0.90 | 2.45e-02 | 2.15e-01 |
| Lachnospiraceae | Carcinoma | Unmatched | 0.43 | 0.21 | 0.90 | 2.52e-02 | 2.15e-01 |
| Bacteroides | Carcinoma | Unmatched | 0.48 | 0.26 | 0.92 | 2.71e-02 | 2.15e-01 |
| Clostridium_sensu_stricto | Carcinoma | Unmatched | 1.89 | 1.04 | 3.43 | 3.79e-02 | 2.80e-01 |
| Barnesiella | Carcinoma | Unmatched | 0.64 | 0.41 | 0.98 | 4.16e-02 | 2.89e-01 |

| Taxon | Tumor | Tissue Group | OR | 95% CI (Lower Bound) | 95% CI (Upper Bound) | P-value | BH |
|---|---|---|---|---|---|---|---|
| Fusobacterium | Carcinoma | Matched | 3.98 | 1.19 | 13.24 | 2.45e-02 | 9.26e-01 |
| Campylobacter | Carcinoma | Matched | 7.80 | 1.20 | 50.88 | 3.18e-02 | 9.26e-01 |
