## Supplementary figures and images for "Leveraging Existing 16S rRNA Gene Surveys to Identify Reproducible Biomarkers in Individuals with Colorectal Tumors"

### Supplementary Materials

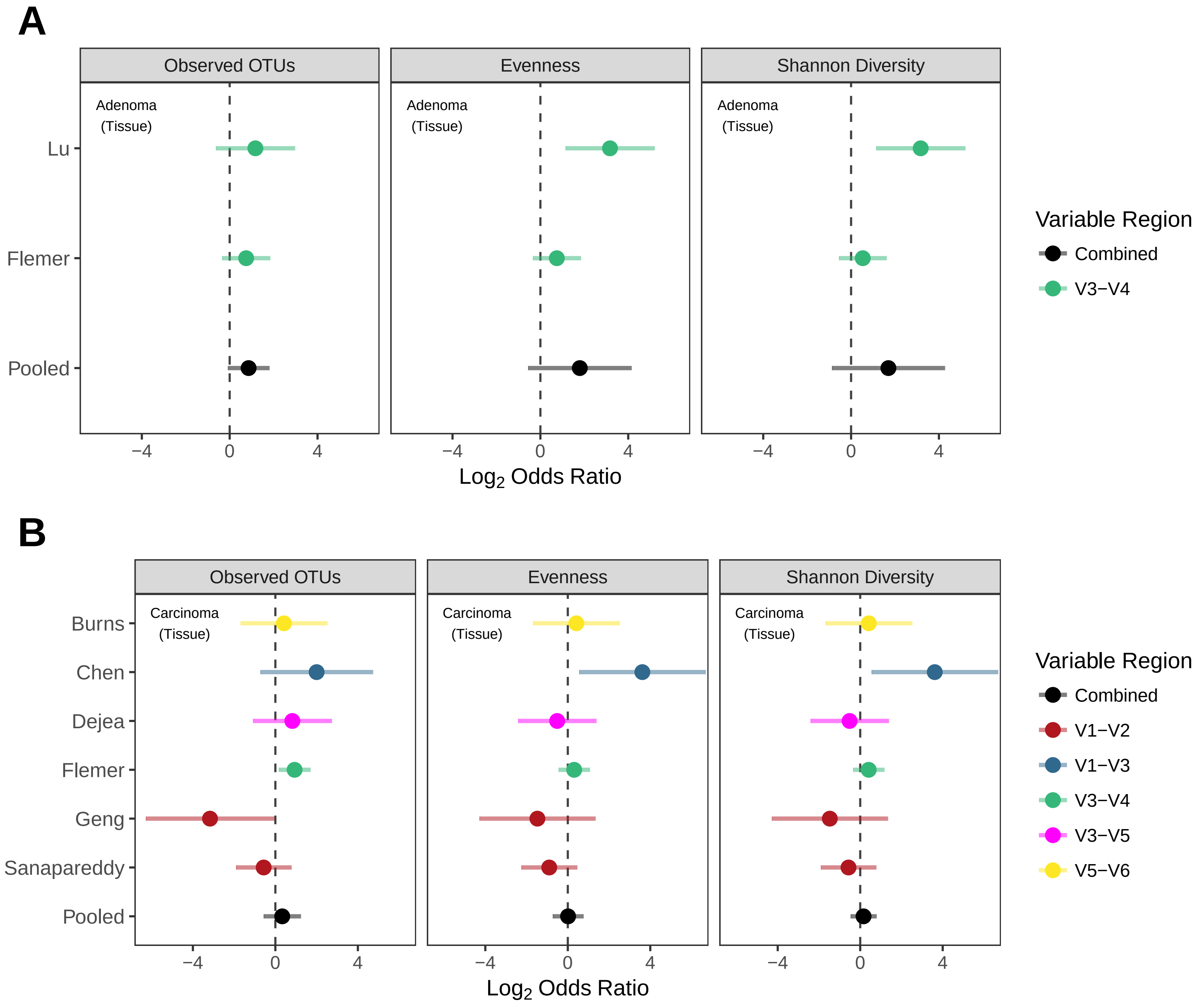

### Supplementary Materials

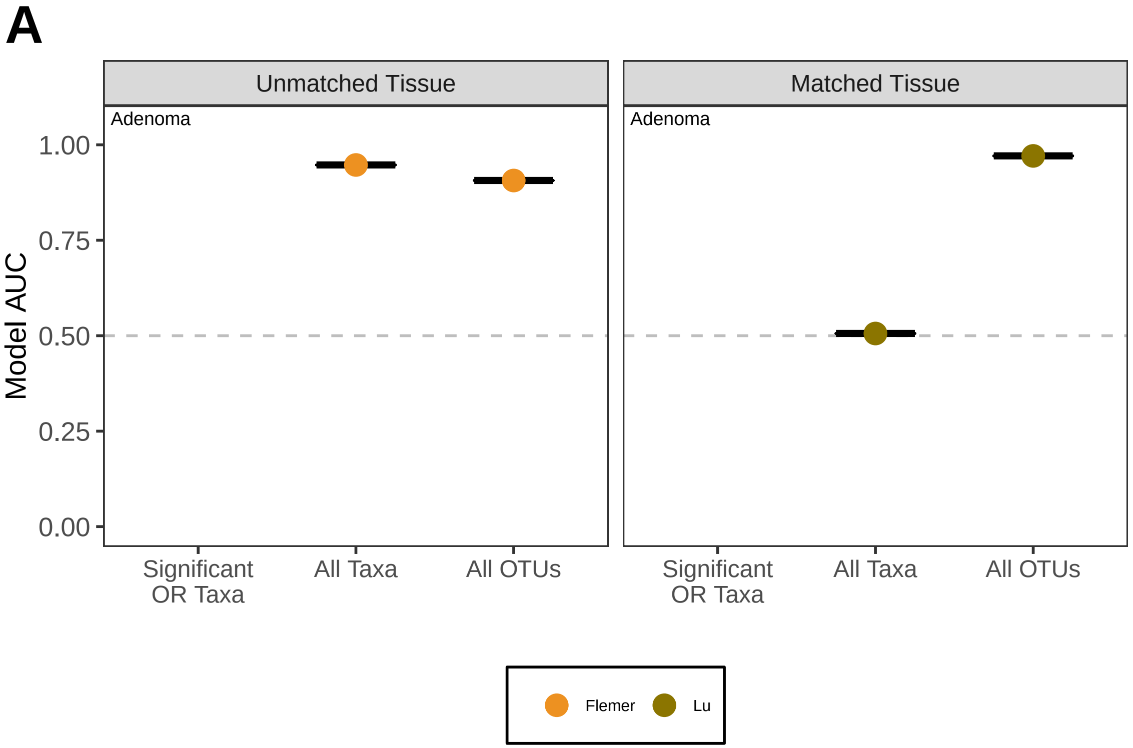

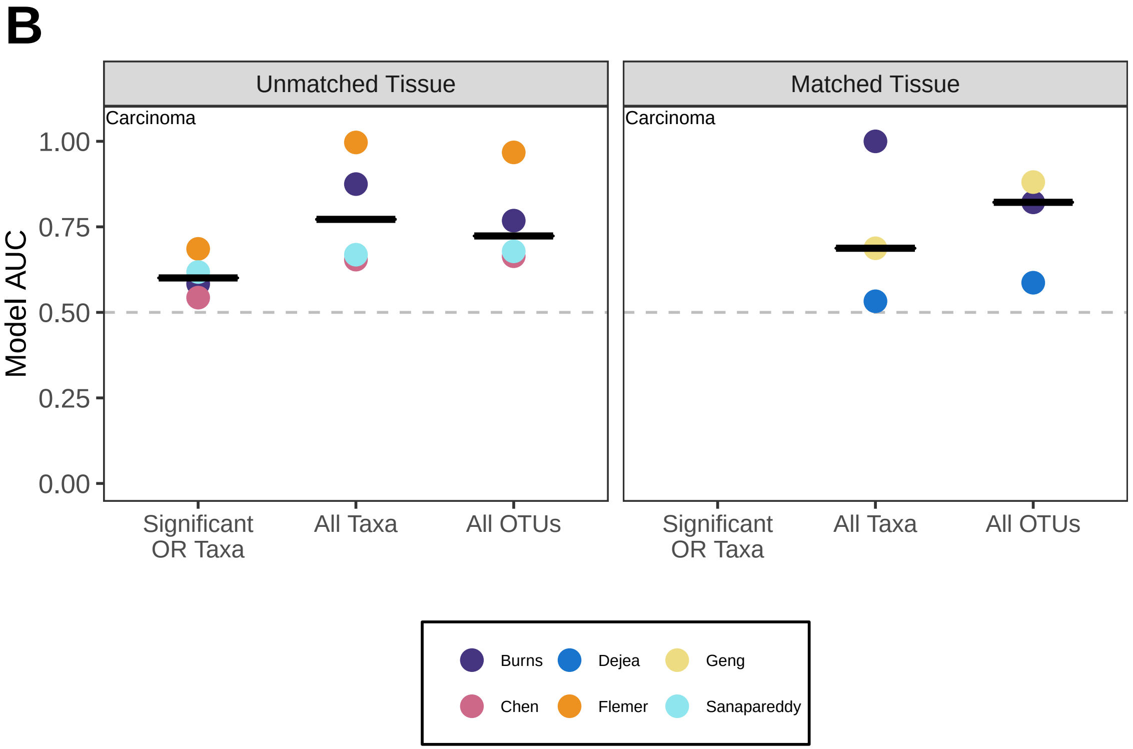

### Supplementary Materials

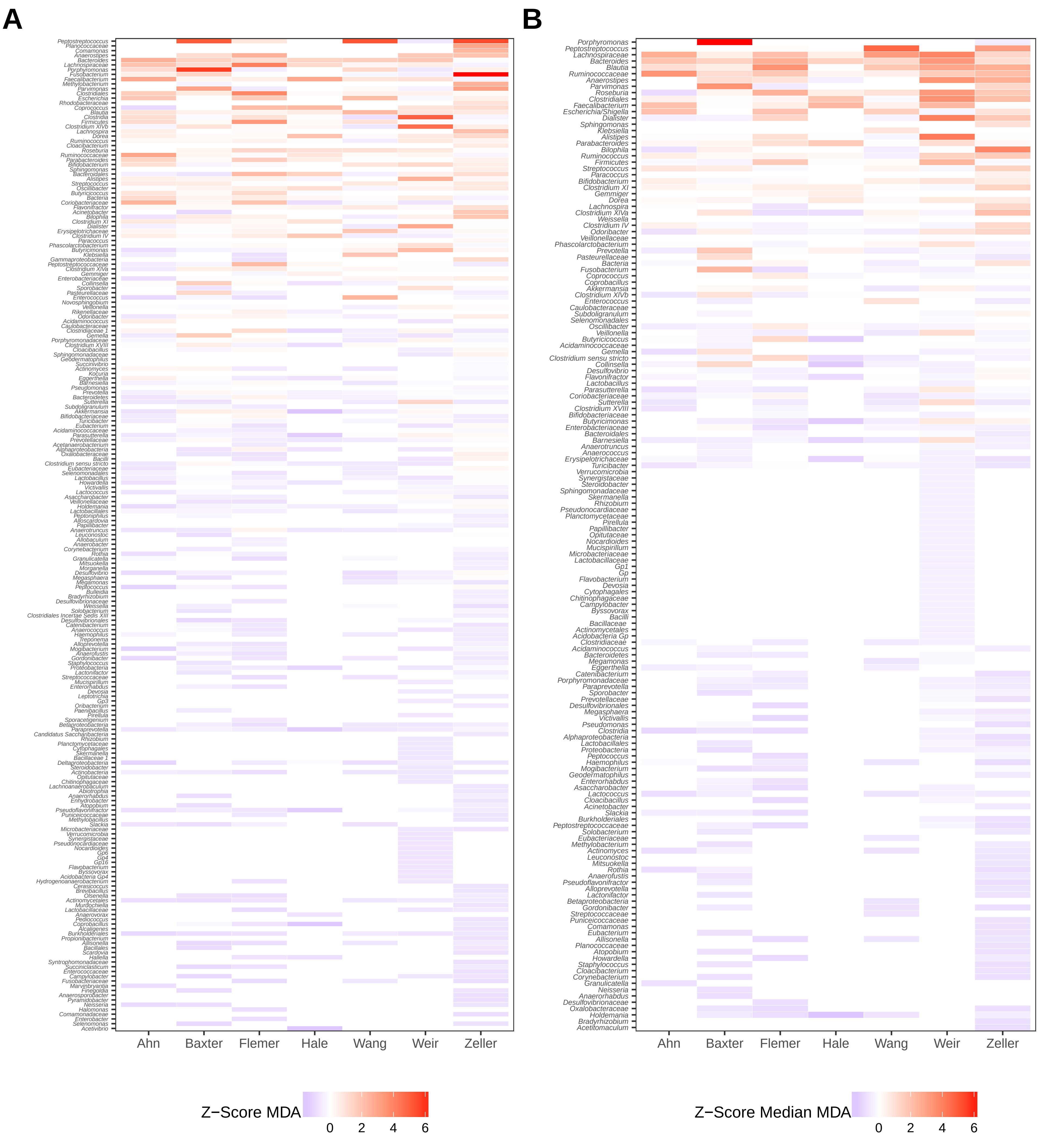

### Supplementary Materials

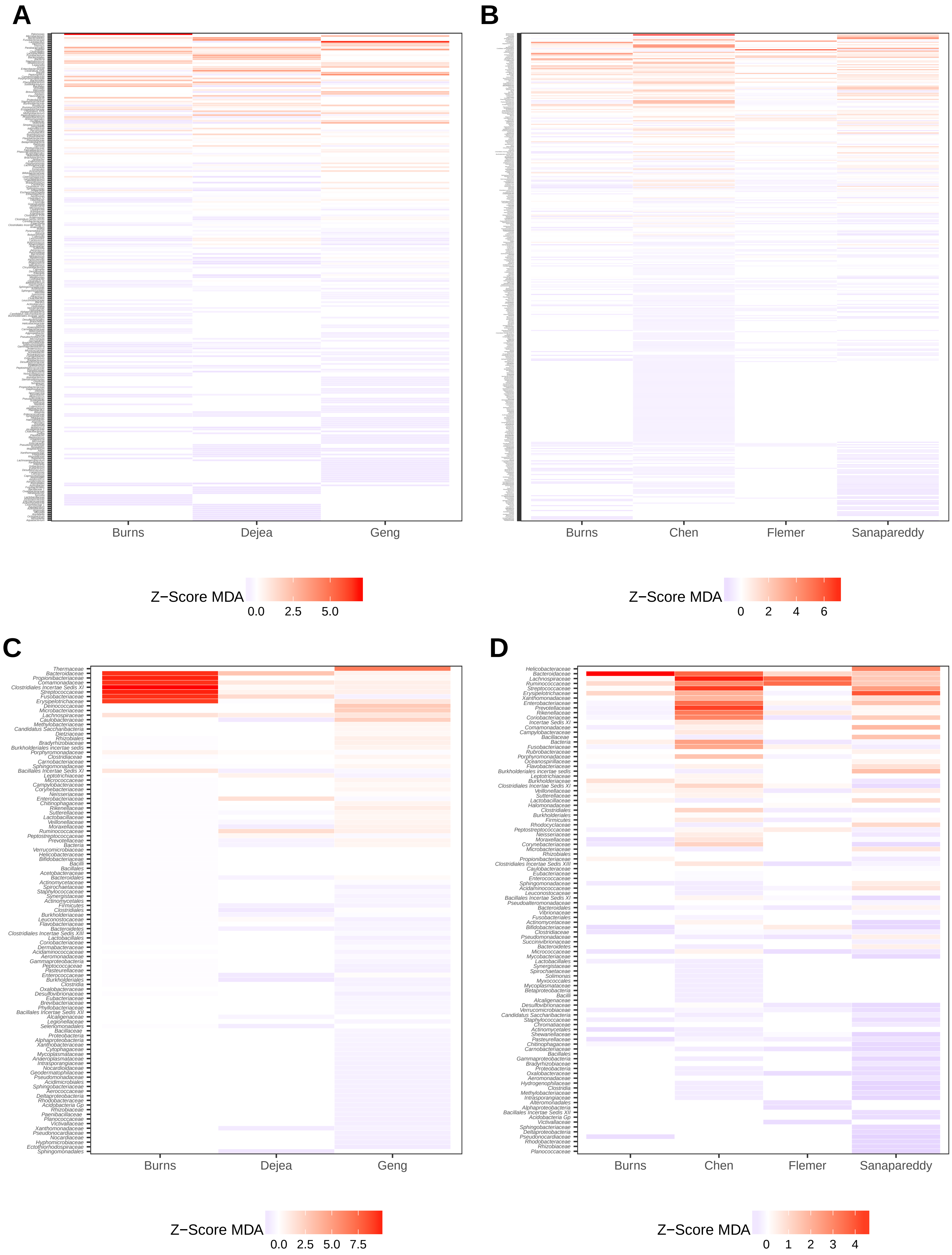

### Supplementary Materials

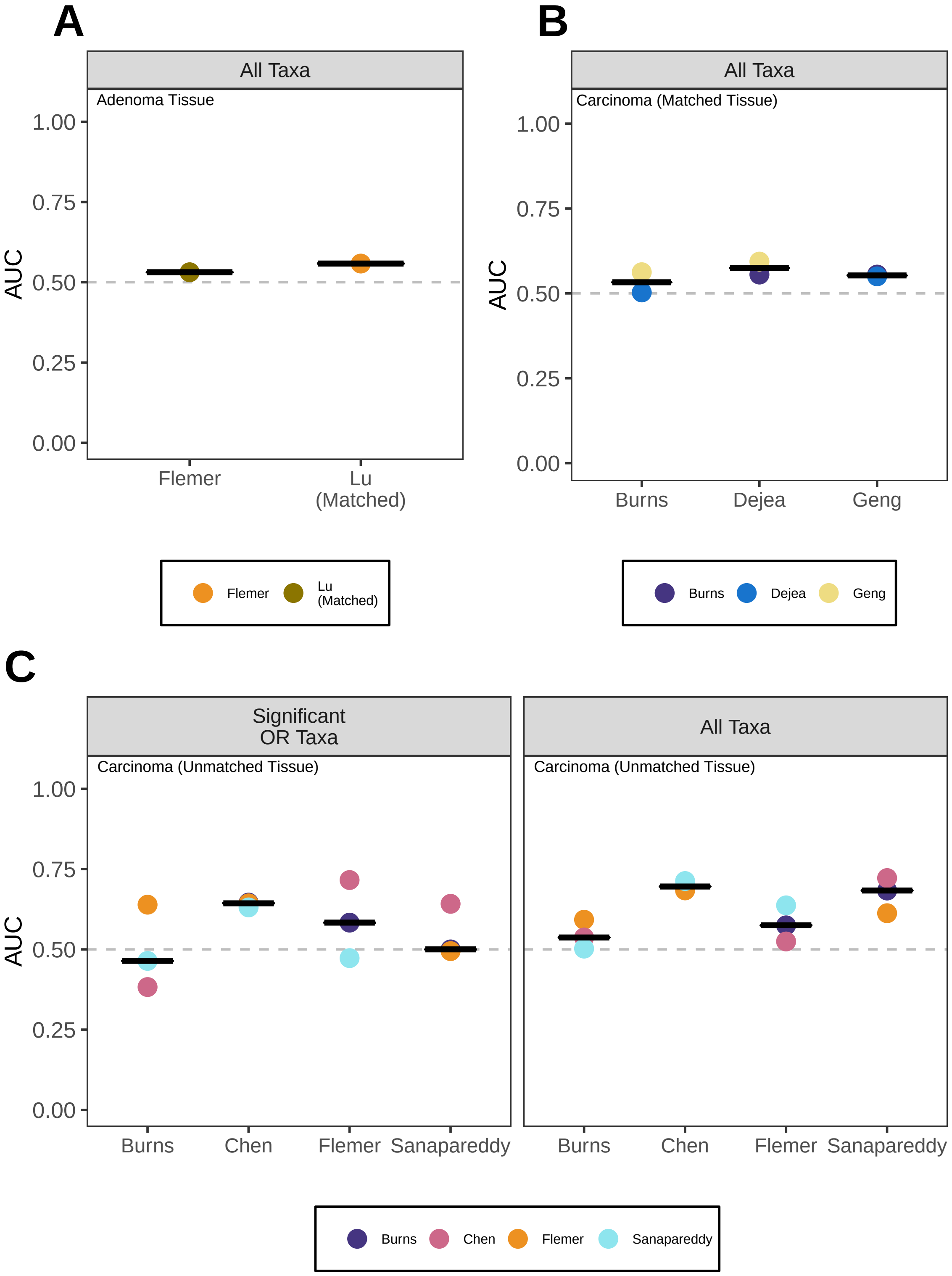
